## Supplemental table and figures for "Benchmarking community-wide estimates of growth potential from metagenomes using codon usage statistics"

August 4, 2022

S5 Cross validation shows MMv2 improves on previous MMv1 model or does just as well. 8
S6 Benchmarking MMv2 against empirically-measured growth rates from the literature. 9
S7 We calculated the mean squared error (MSE) of the predicted median minimum doubling time of mixtures of genomes to the median prediction for individual genomes. 10
S8 MMV2 predictions have somewhat lower error for low-$\Psi$ ($< 0.6$) samples. . . . . . 11
S9 Temperature correction is important for RefSeq genome mixtures with simulated sample temperatures. . . . . . . . . . . . . . . . . . . . . . . . . . . . . . . . . . 12
S10 Temperature correction is important for human gut genome mixtures with simulated sample temperatures. . . . . . . . . . . . . . . . . . . . . . . . . . . . . . . . . 13
S11 Temperature correction is important for ocean surface genome mixtures with simulated sample temperatures. . . . . . . . . . . . . . . . . . . . . . . . . . . . . . 14
S12 MMv2 has longer run times than MMv1 for HMP metagenomes when $\Psi > 0.6$. . . . 15
S13 Short genes produce biased CUB estimates. We took the set of all genes in a genome at least 510 nucleotides long and truncated these genes to progressively shorter lengths. 16
S14 Short ribosomal proteins genes produce biased CUB estimates. . . . . . . . . . . . . 17
S15 Codon usage biases from the synthetic metagenomes discussed in Fig 4. . . . . . . . 18
S16 Predicted maximum growth rates from genome mixtures and real metagenomes are stable under subsampling as long as at least 5% of genes are sampled. . . . . . . . . 19
S17 Predicted maximum growth rates from genome mixtures are accurate under subsampling as long as at least 5% of genes are sampled. . . . . . . . . . . . . . . . . . . . . 20

S18 Predicted maximum growth rates (unweighted mode, no coverage information) from genome mixtures and real metagenomes are stable under subsampling as long as at least 5% of genes are sampled. . . . . . . . . . . . . . . . . . . . . . . . . . . . . . 21
S19 Predicted maximum growth rates (unweighted mode, no coverage information) from genome mixtures are accurate under subsampling as long as at least 5% of genes are sampled. . . . . . . . . . . . . . . . . . . . . . . . . . . . . . . . . . . . . . . . . . . 22
S20 Max. growth rate predictions capture growth rate variation across single-genus species mixtures. . . . . . . . . . . . . . . . . . . . . . . . . . . . . . . . . . . . . . . 23
S21 The results of Fig 4 are unchanged when sampling a larger number (10%) of genes predicted from reads for each sample. . . . . . . . . . . . . . . . . . . . . . . . . . . 24

Table S1: **Maximum growth rate models.** *MMv2 is a piece-wise combination of MMBC and MMv1, where MMv1 is used if $\Psi < 0.6$ and MMBC is used otherwise

| Model | CUB Metrics | Other Variables | CUB Calculated at the Single-Gene Level |
|---|---|---|---|
| MMBC* | $\Delta$ MILC | GC, OGT | Y |
| MMv1* | $MILC_{HE}$ | OGT | N |
| growthpred | $\Delta$ENC', S | OGT | Y |
| gRodon "full" | $MILC_{HE}$ | $\Psi$, Codon Pair Bias, OGT | N |

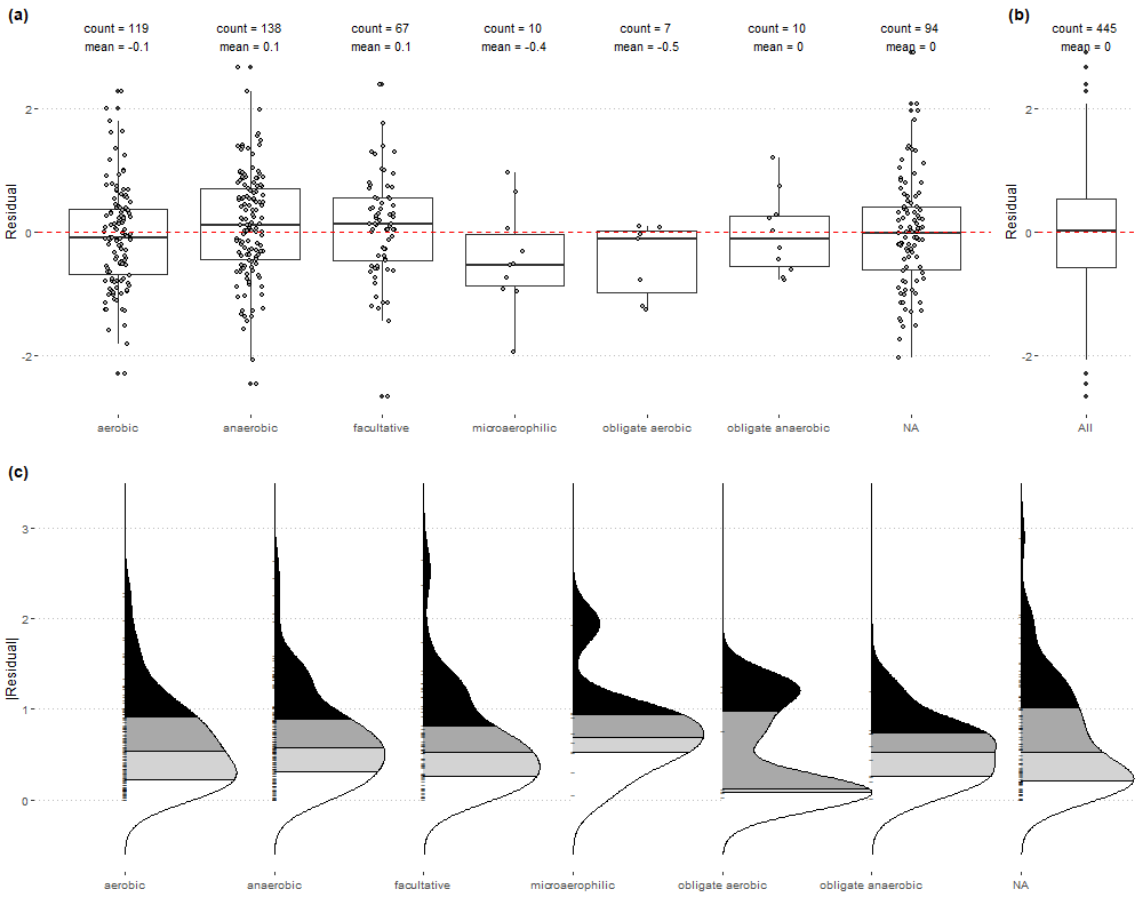

Figure S1: Predictions from gRodon are not impacted by the metabolic oxygen use of an organism. (a) Metabolic use had no significant effect on model residuals (ANOVA, $p = 0.077$), nor (b) on the absolute value of the model residuals (ANOVA, $p = 0.79$) for gRodon's "full" mode.

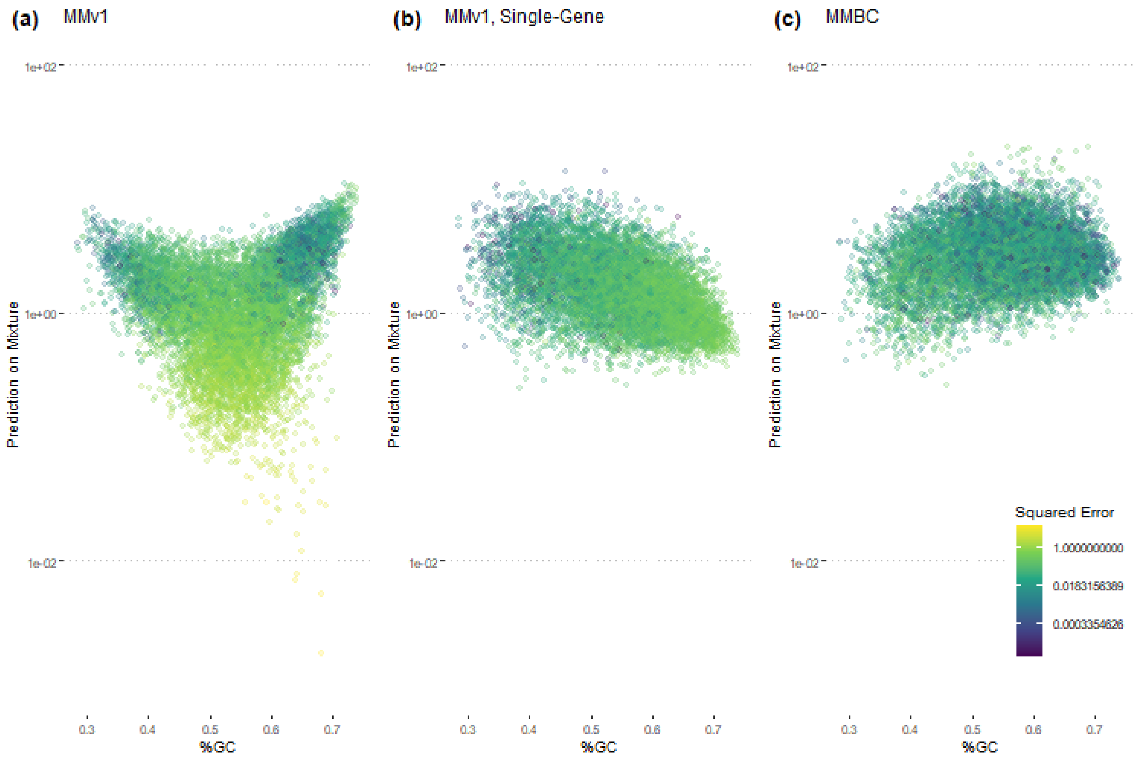

Figure S2: Bias-corrected metagenome mode (MMBC) corrects the strong GC bias seen in metagenome mode V1 (MMv1). Using a per-gene estimation of the background codon usage with the MMv1 model (see Methods) only partially corrects for this pattern. By normalizing CUB estimates and adding an explicit GC correction (see Methods), MMBC is able to effectively remove this GC bias. Each point in these plots represents a simulated 2-organism mixed community drawn from our set of RefSeq genomes.

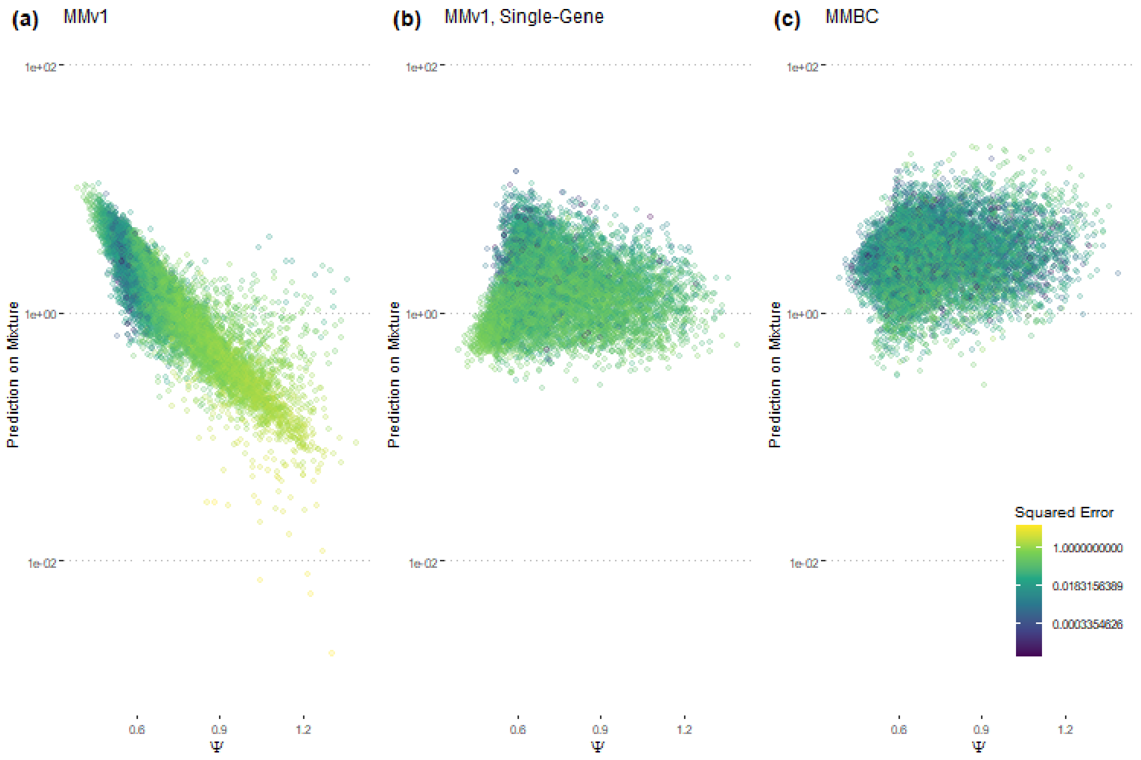

Figure S3: Bias-corrected metagenome mode (MMBC) corrects the strong Ψ bias seen in metagenome mode V1 (MMv1). Using a per-gene estimation of the background codon usage with the MMv1 model (see Methods) mostly corrects for this pattern. By normalizing CUB estimates and adding an explicit GC correction (see Methods), MMBC is able to further mitigate this Ψ bias. Each point in these plots represents a simulated 2-organism mixed community drawn from our set of RefSeq genomes.

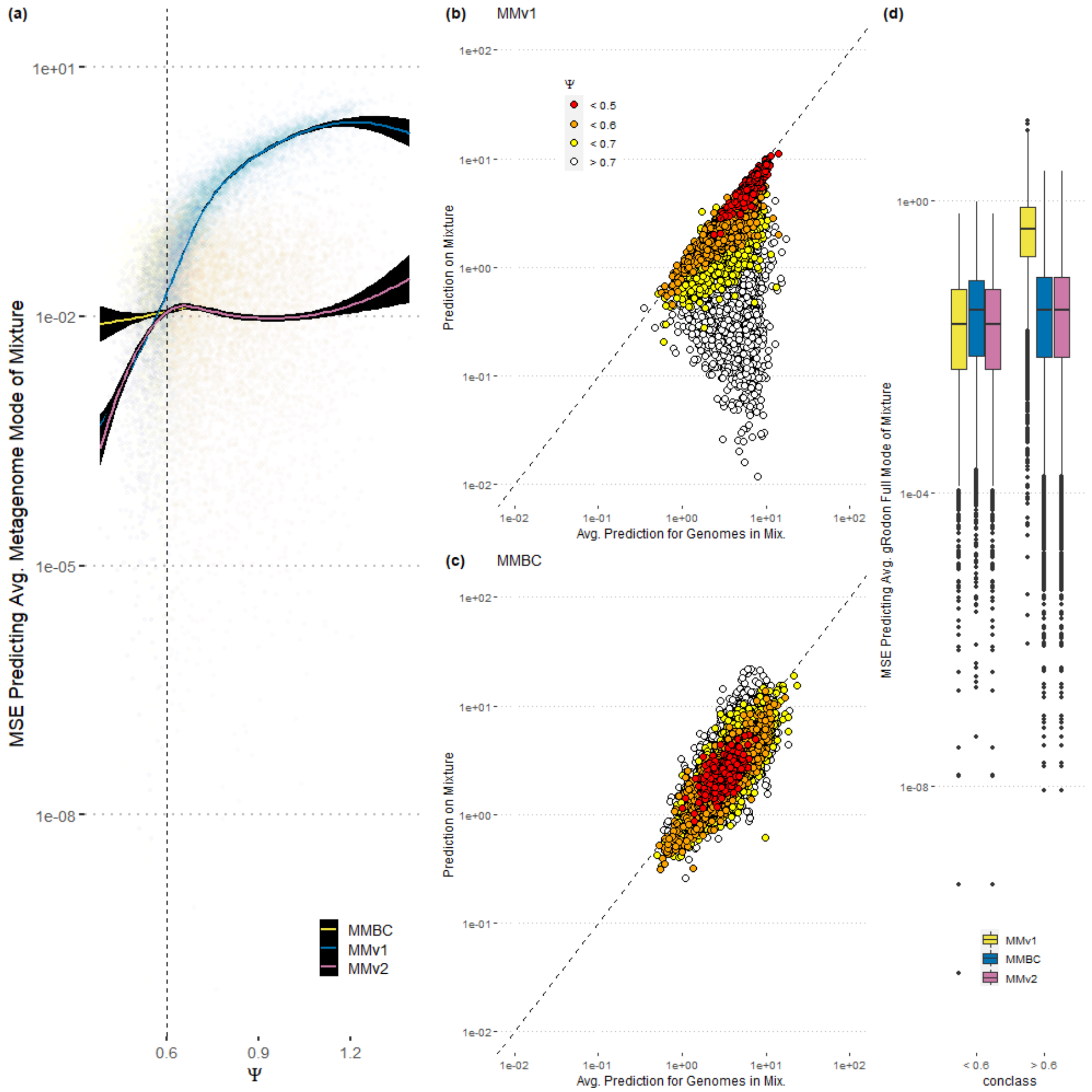

Figure S4: By combining bias-corrected metagenome mode (MMBC) and metagenome mode v1 (MMv1) we are able to construct a superior predictor (MMv2). (a-c) These panel show how well each predictor converges to "itself". That is, if we predict the max. growth rate of each genome in a mixture using that model and take the average, how well does the prediction on the mixed community estimate that value? For low Ψ values, MMv1 outperforms MMBC in terms of this convergence (see discussion of bias-variance tradeoff in main text). MMv2 switches between these two models based on a Ψ threshold of 0.6. Each point in these plots represents a simulated 2-organism mixed community drawn from our set of RefSeq genomes. (d) This panel shows how well each predictor is at estimating the "real" community-wide average growth rates. That is, if we predict the max. growth rate of each genome in a mixture using gRodons "full" mode (the best predictor we have for single genomes) and take the average, how well does the prediction on the mixed community estimate that value?

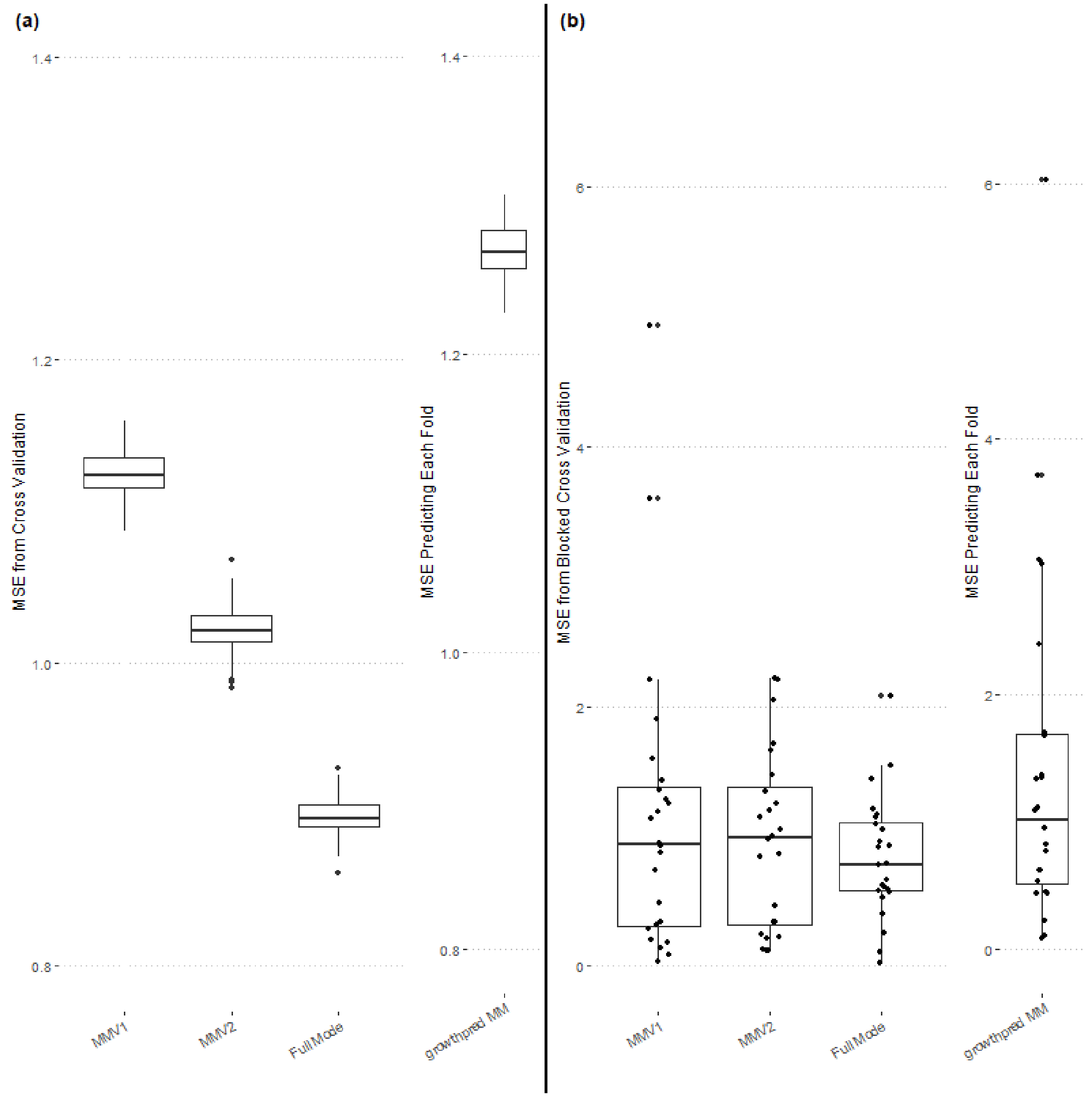

Figure S5: Cross validation shows MMv2 improves on previous MMv1 model or does just as well. (a) Mean squared error (MSE) from random cross validation of each model on the training data. (b) Blocked cross validation of model fit using phyla as folds. Because growthpred was trained on a different dataset it's data, the error reported for this model is not technically cross validated, but given that the training set used for growthpred is highly overlapping with the training data used here this may give growthpred a slight advantage (particularly in blocked cross validation where only the growthpred model is trained on species from the phylum in the held out folds).

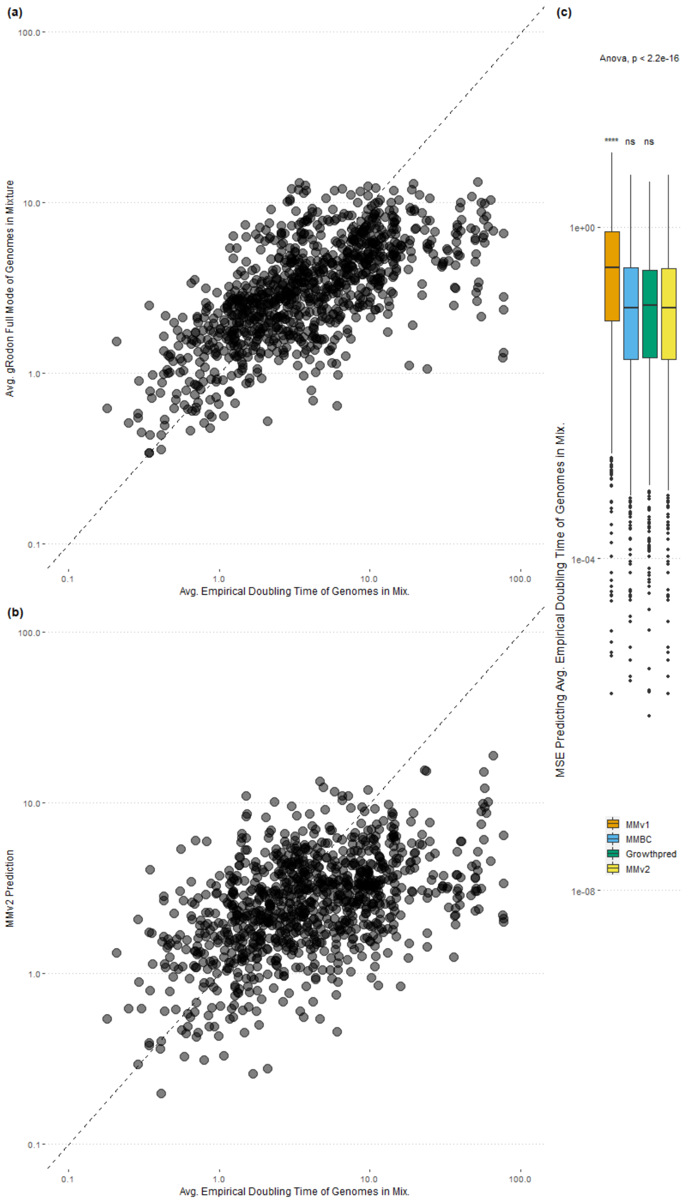

Figure S6: Benchmarking MMv2 against empirically-measured growth rates from the literature. (a) Full mode estimates used as a benchmark recapitulate measured estimates. (b,c) MMv2 does a comparable job when benchmarked against empirically measured rates to when benchmarked against inferred rates (Fig 1) and consistently improves on MMv1 predictions.

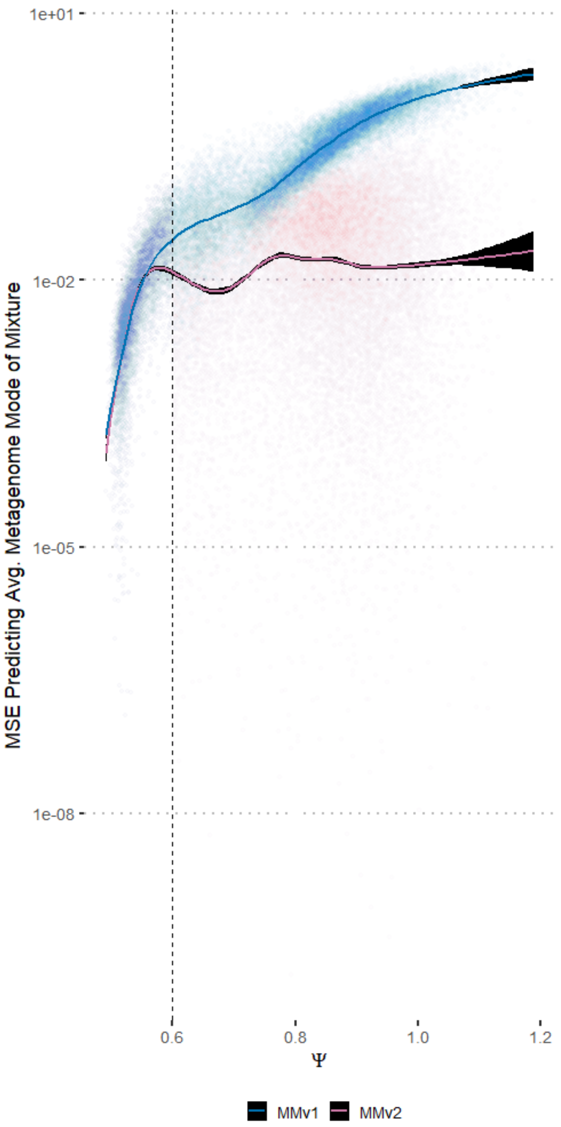

Figure S7: We calculated the mean squared error (MSE) of the predicted median minimum doubling time of mixtures of genomes to the median prediction for individual genomes. Results shown for all genome mixtures ($n = 30,000$; drawn from RefSeq, human gut isolates, and marine SAGs). $\Psi$ is a measure of codon-usage dissimilarity across highly-expressed genes in a sample, and low values of $\Psi$ indicate high similarity (see Results and Discussion).

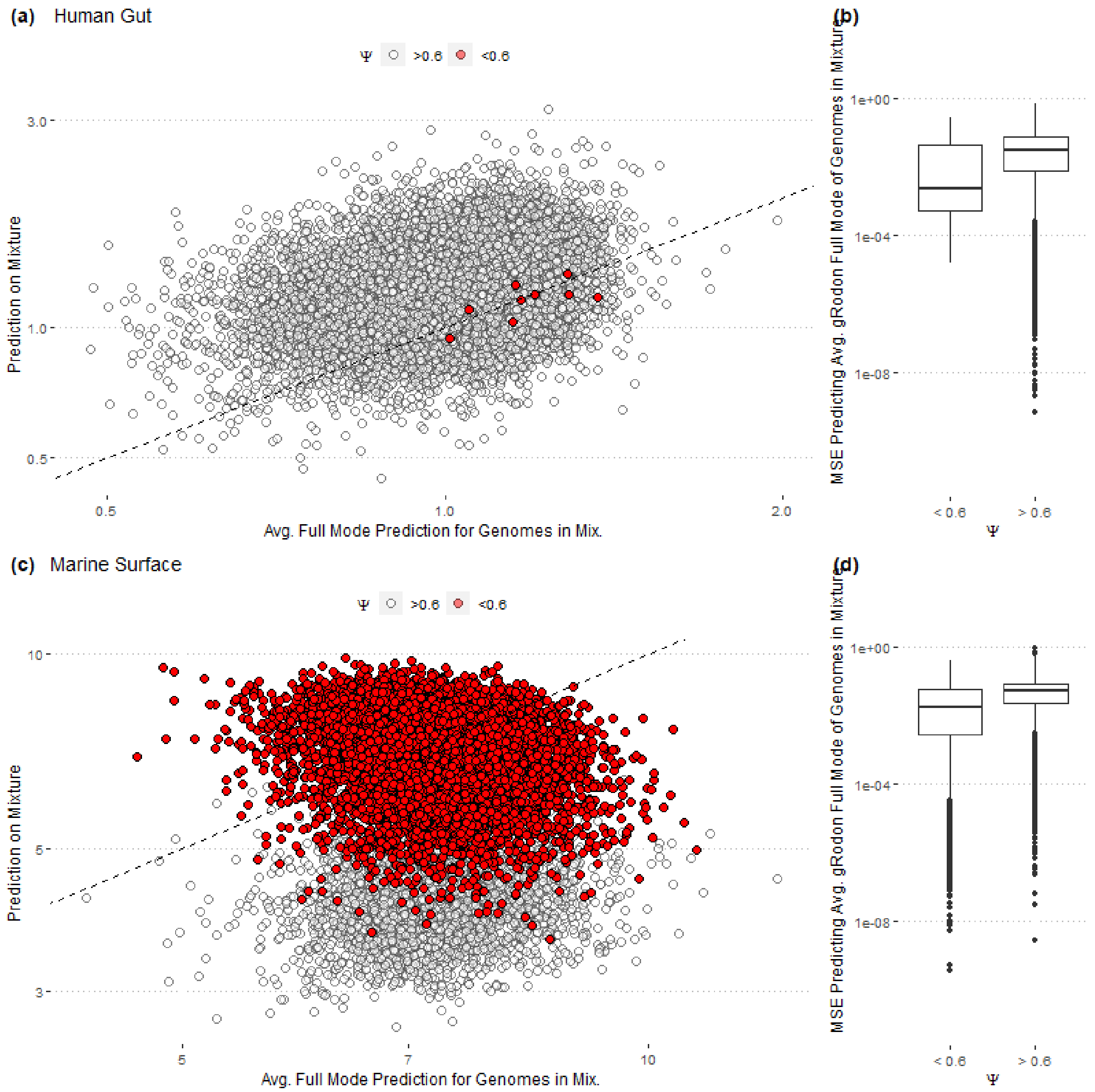

Figure S8: MMV2 predictions have somewhat lower error for low-Ψ ($< 0.6$) samples. RefSeq mixtures not shown because no RefSeq mixtures had $\Psi < 0.6$.

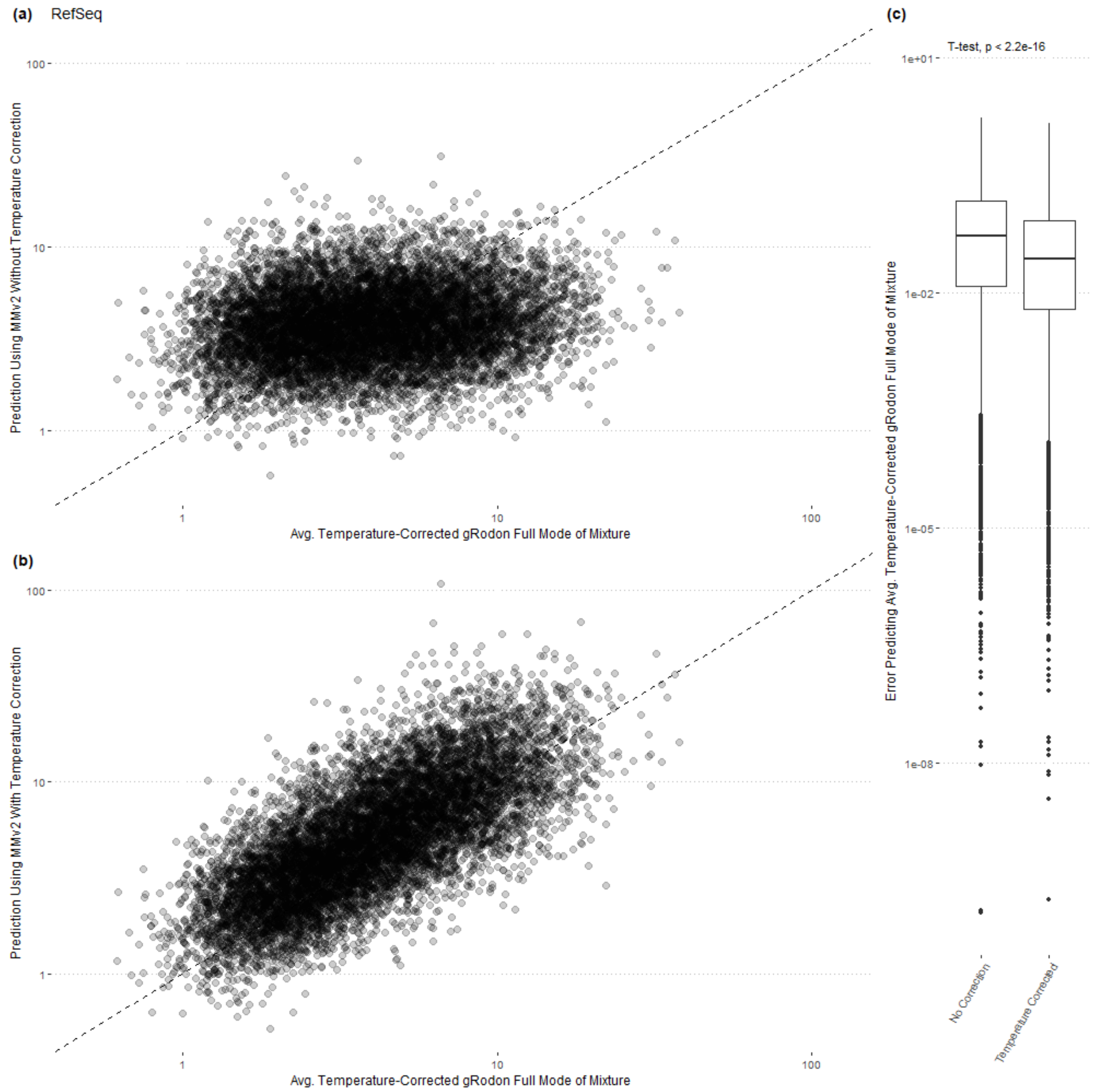

Figure S9: Temperature correction is important for RefSeq genome mixtures with simulated sample temperatures. For each genome mixture, a sample temperature between 0 and 60°C was drawn from a uniform distribution and the optimal growth temperature of each species in the mixture was taken as the sum of this sample-wide value and a draw from a normal distribution with mean zero and standard deviation of 10. These organism-level temperatures were used to predict the individual growth rates of genomes (x-axis), and the sample temperature was used to predict the community level growth rate in panel (b). This approach was used to account for possible variation in an organism's OGT relative to the conditions it may be found in.

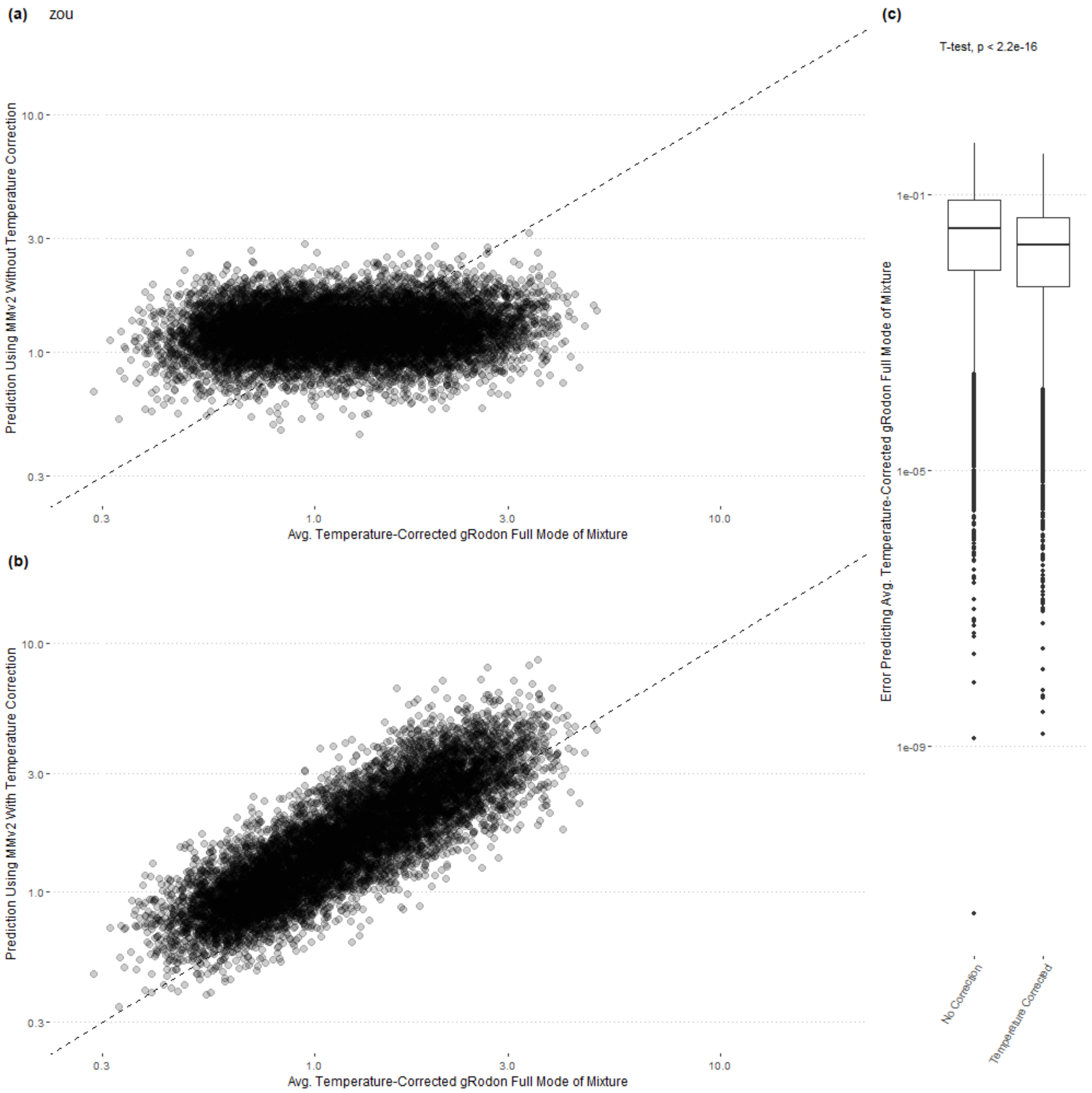

Figure S10: Temperature correction is important for human gut genome mixtures with simulated sample temperatures. For each genome mixture, a sample temperature between 0 and 60°C was drawn from a uniform distribution and the optimal growth temperature of each species in the mixture was taken as the sum of this sample-wide value and a draw from a normal distribution with mean zero and standard deviation of 10. These organism-level temperatures were used to predict the individual growth rates of genomes (x-axis), and the sample temperature was used to predict the community level growth rate in panel (b). This approach was used to account for possible variation in an organism's OGT relative to the conditions it may be found in.

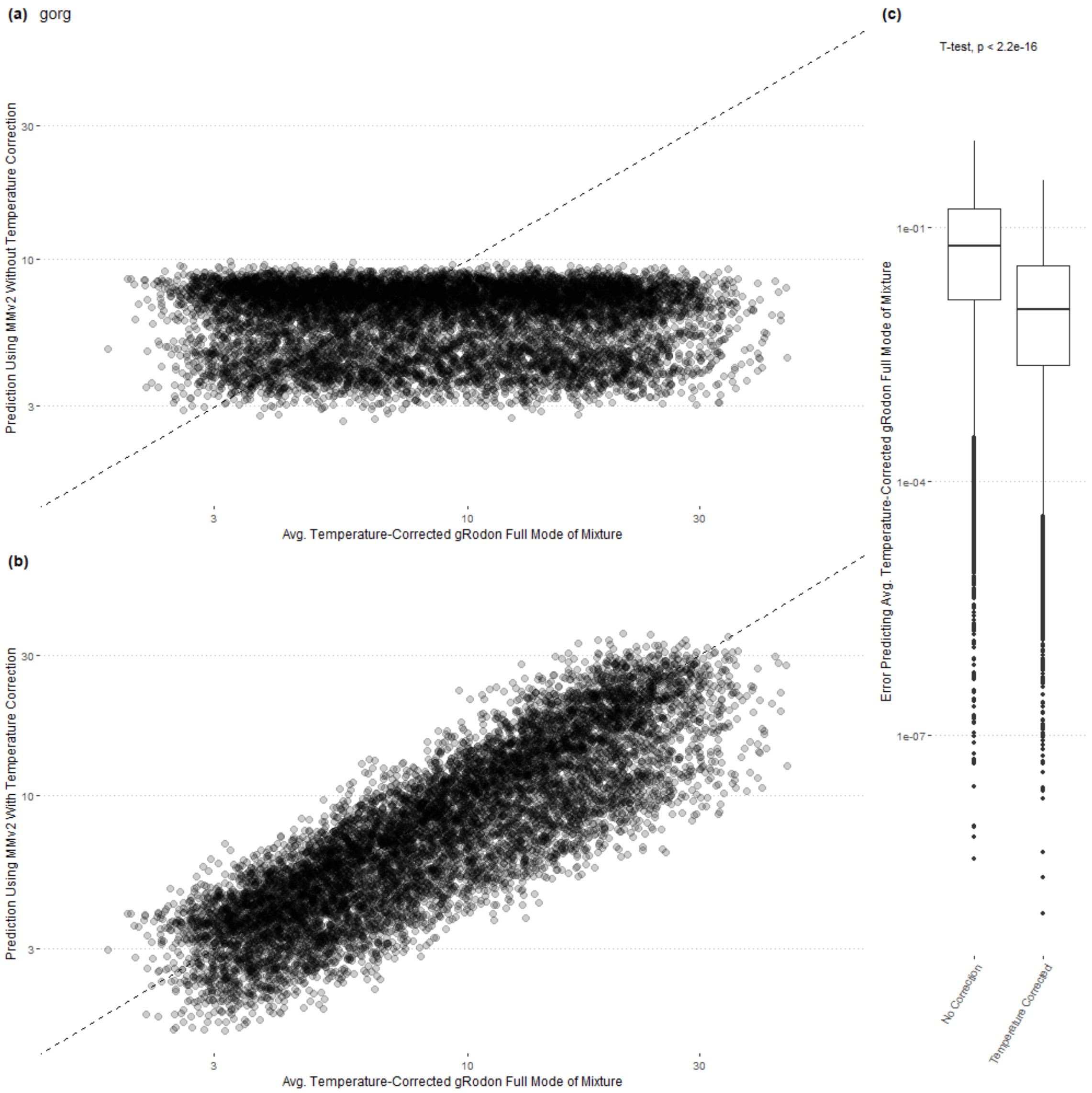

Figure S11: Temperature correction is important for ocean surface genome mixtures with simulated sample temperatures. For each genome mixture, a sample temperature between 0 and 60°C was drawn from a uniform distribution and the optimal growth temperature of each species in the mixture was taken as the sum of this sample-wide value and a draw from a normal distribution with mean zero and standard deviation of 10. These organism-level temperatures were used to predict the individual growth rates of genomes (x-axis), and the sample temperature was used to predict the community level growth rate in panel (b). This approach was used to account for possible variation in an organism's OGT relative to the conditions it may be found in.

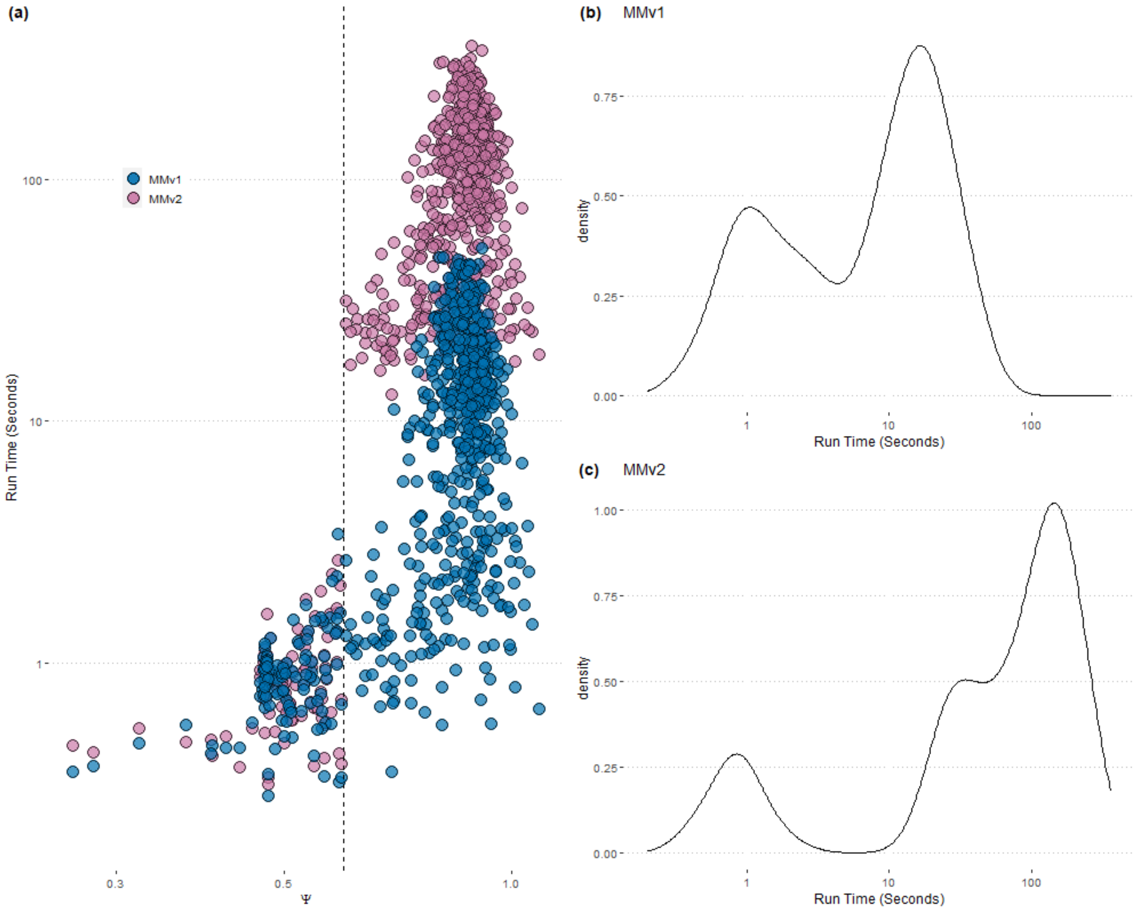

Figure S12: MMv2 has longer run times than MMv1 for HMP metagenomes when $\Psi > 0.6$.

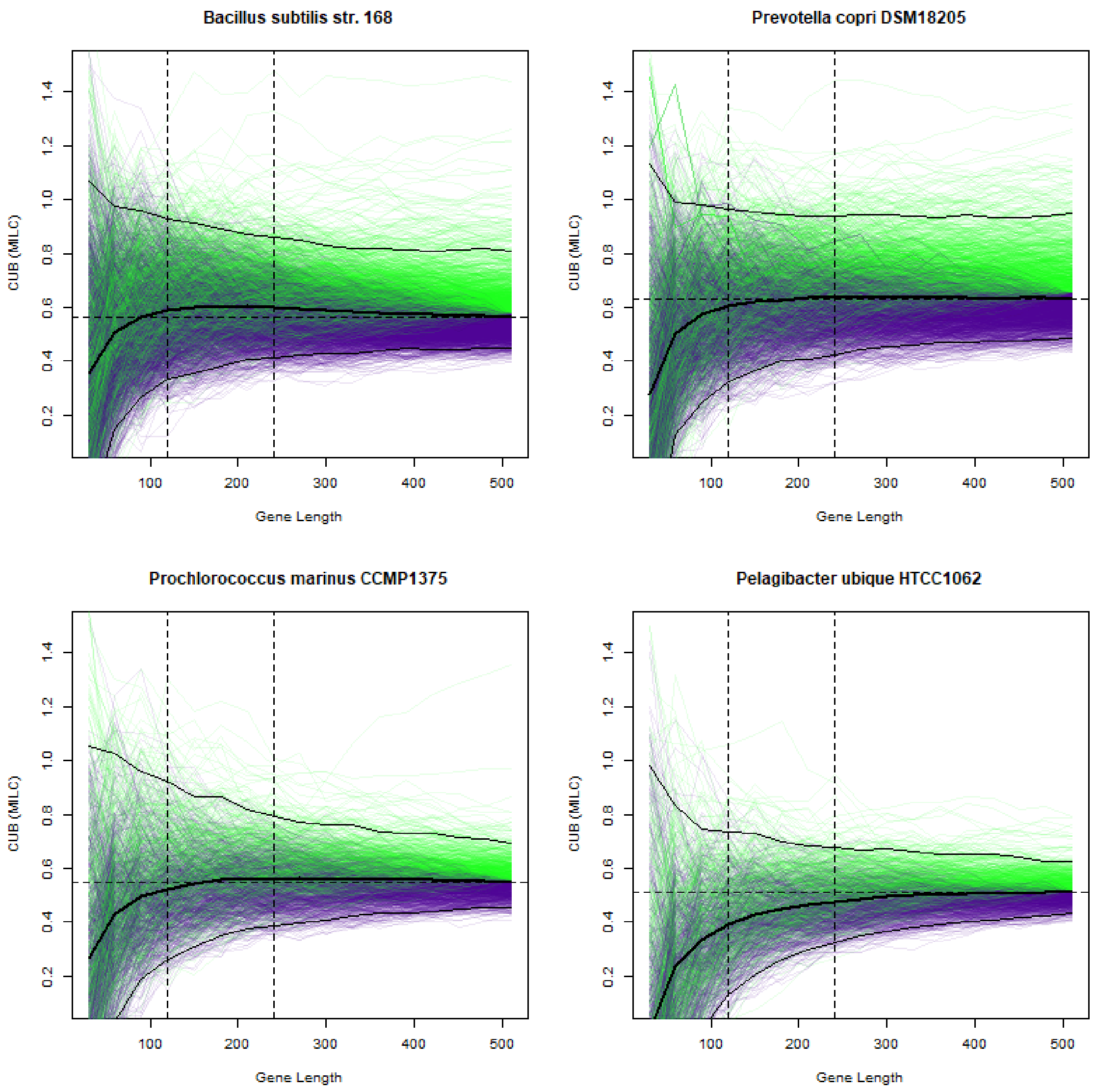

Figure S13: Short genes produce biased CUB estimates. We took the set of all genes in a genome at least 510 nucleotides long and truncated these genes to progressively shorter lengths. Horizontal dashed line is the mean CUB of all genes at length 510 and we have colored individual gene tracts based on whether they start above (green) or below (purple) that value. Vertical dashed lines at 240bp and 120bp for reference. Thick solid black line is the median CUB for each truncation length and thin solid black lines represent the 97.5% and 2.5% percentiles at each truncation length. The species in the top two panels are fast growing organisms that can be found at least sometimes in the human gut, and the species in the bottom two panels are two slow-growing marine organisms.

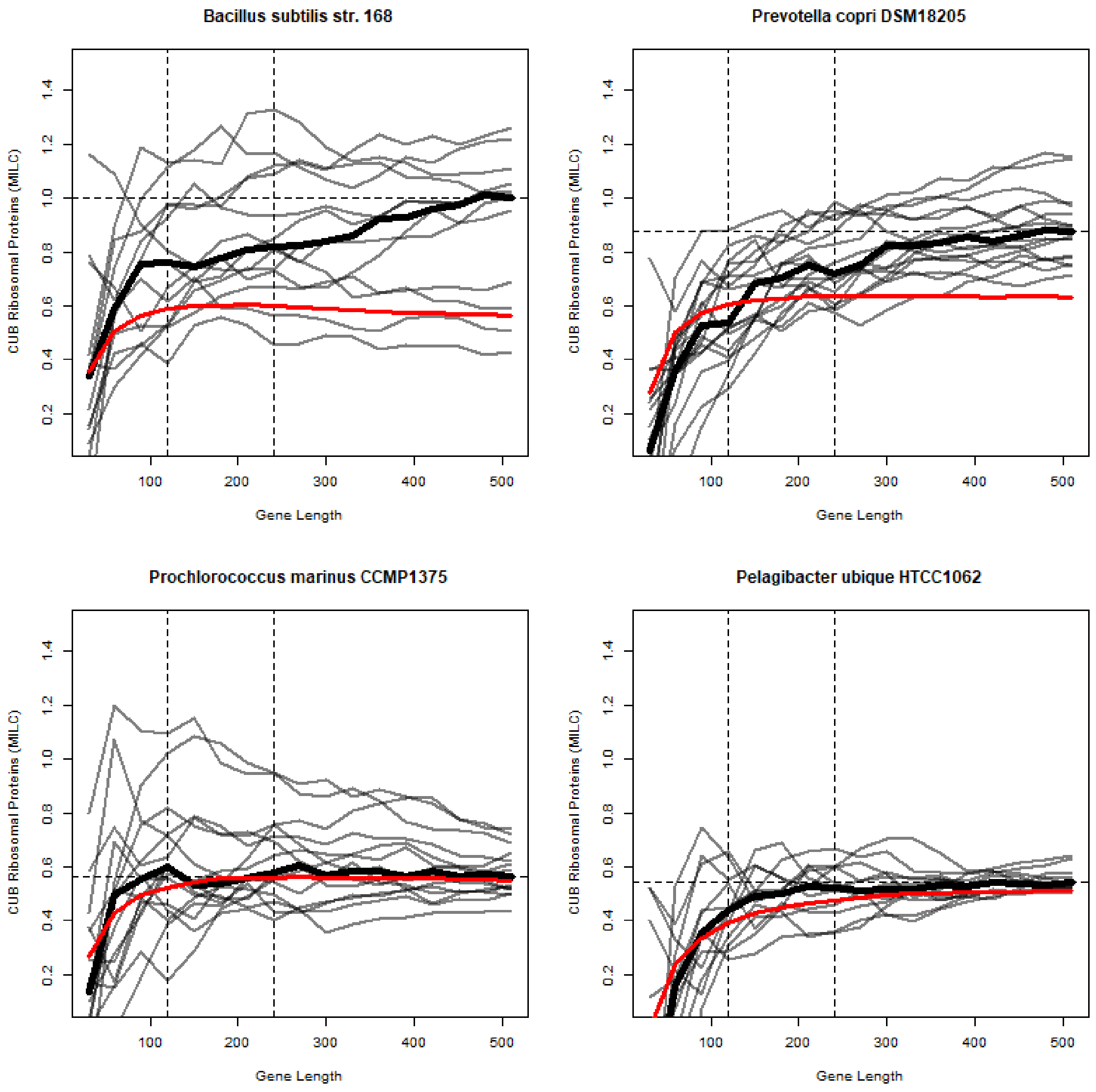

Figure S14: Short ribosomal proteins genes produce biased CUB estimates. We took the set of genes coding for ribosomal proteins in a genome at least 510 nucleotides long and truncated these genes to progressively shorter lengths. Horizontal dashed line is the mean CUB of all ribosomal protein genes at length 510 gray lines are individual gene tracts. Vertical dashed lines at 240bp and 120bp for reference. Thick solid black line is the median CUB of the ribosomal proteins for each truncation length and thick solid red line is the median CUB of all genes in a genome for each truncation length for reference (as in S5 Fig). The species in the top two panels are fast growing organisms that can be found at least sometimes in the human gut, and the species in the bottom two panels are two slow-growing marine organisms.

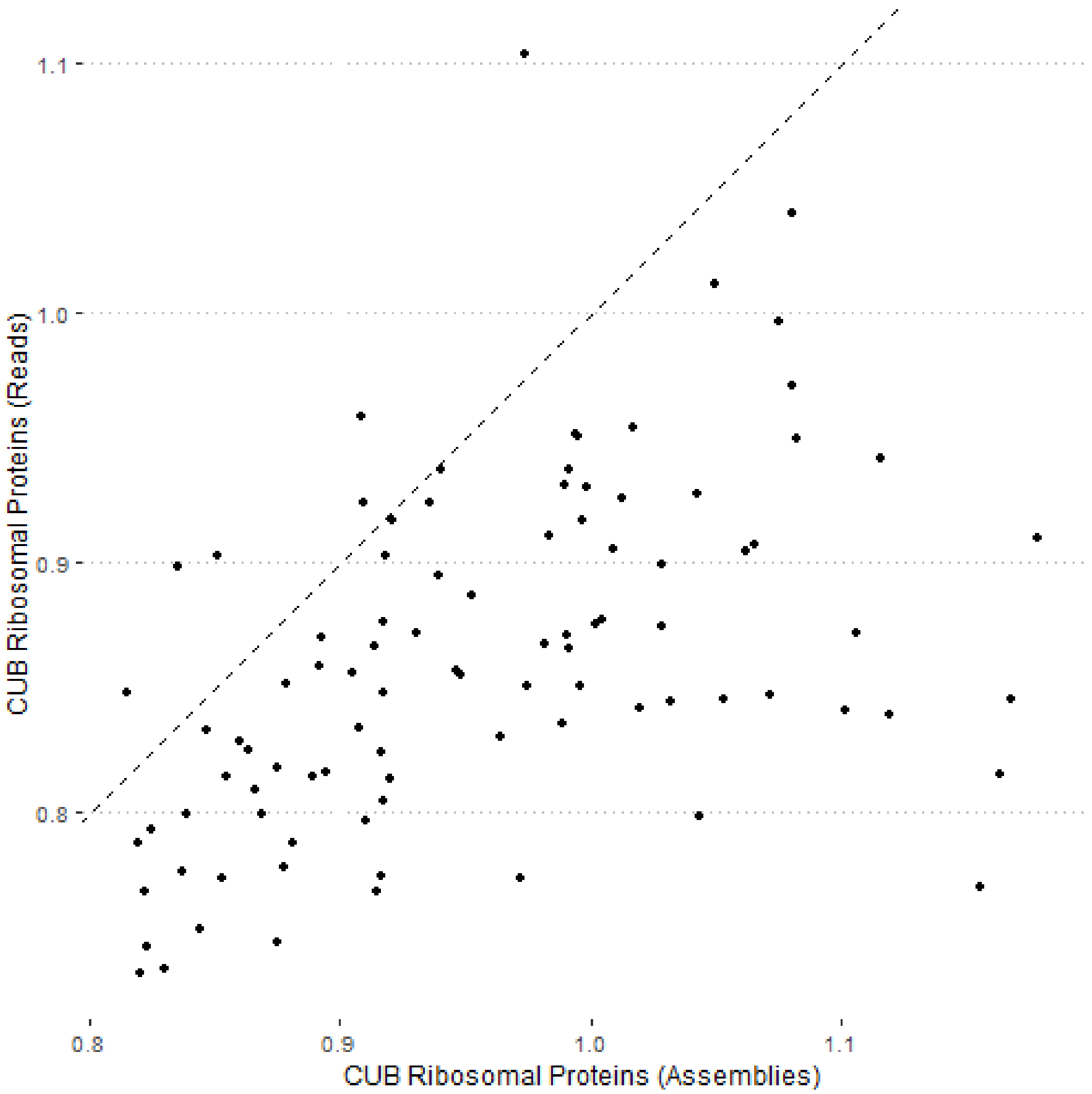

Figure S15: Codon usage biases from the synthetic metagenomes discussed in Fig 4.

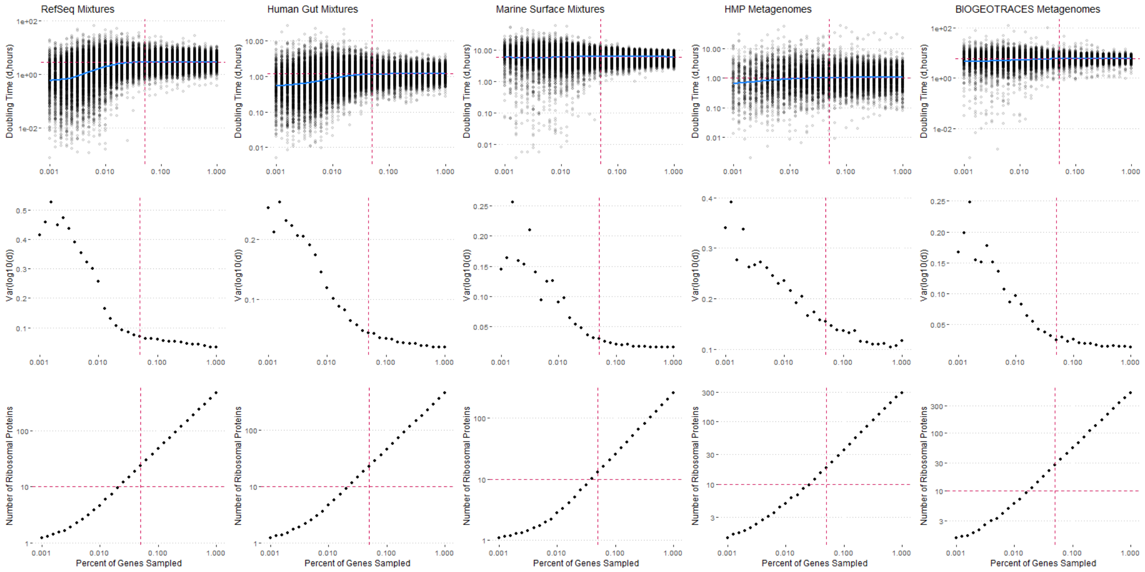

Figure S16: Predicted maximum growth rates from genome mixtures and real metagenomes are stable under subsampling as long as at least 5% of genes are sampled. After this threshold doubling time estimates appear to be slightly biased towards faster growth, and the variance of the estimated minimum doubling time increases dramatically. This 5% threshold corresponds to a subsampled metagenome that still has at least 30-50 ribosomal protein genes sampled. Vertical dashed line denoted the 5% cutoff. Horizontal dashed line denotes a cutoff of 10 ribosomal proteins that is typically used for individual genomes and appears to be too low for metagenomes (30-50 is a better cutoff).

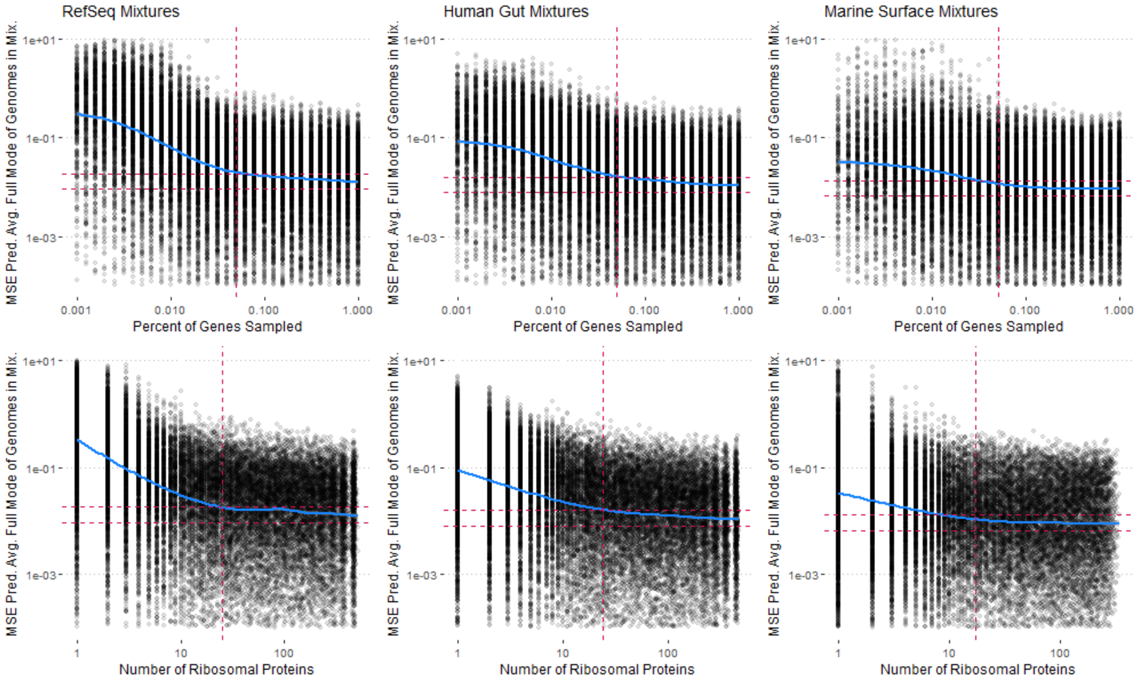

Figure S17: Predicted maximum growth rates from genome mixtures are accurate under subsampling as long as at least 5% of genes are sampled. After this threshold errors increase dramatically. Vertical dashed line denoted the 5% cutoff. Horizontal dashed lines denotes mean error of prediction from non-subsampled mixtures and that error value doubled.

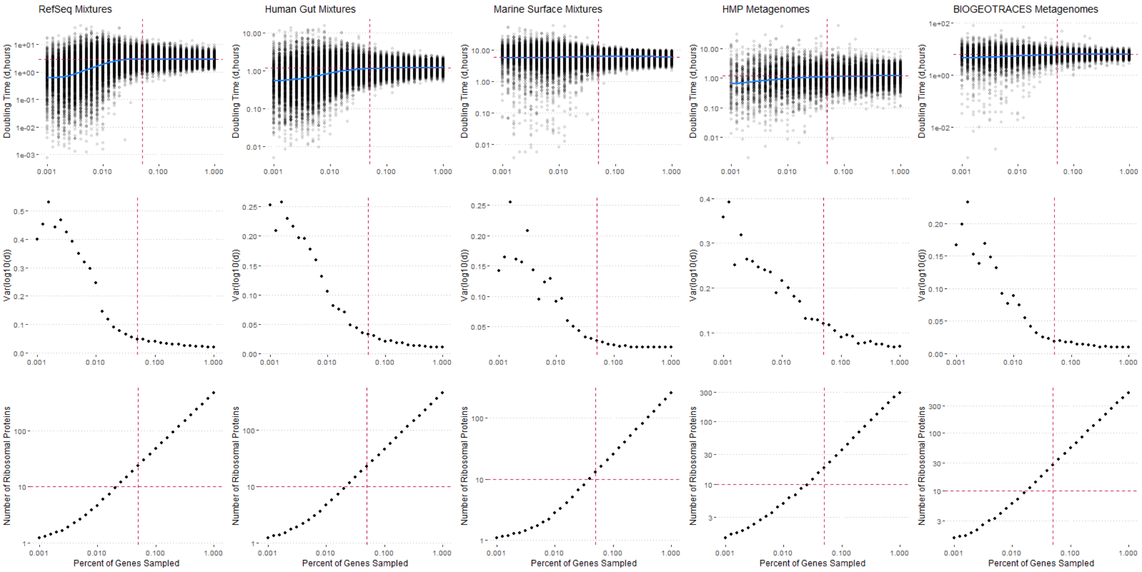

Figure S18: Predicted maximum growth rates (unweighted mode, no coverage information) from genome mixtures and real metagenomes are stable under subsampling as long as at least 5% of genes are sampled. After this threshold doubling time estimates appear to be slightly biased towards faster growth, and the variance of the estimated minimum doubling time increases dramatically. This 5% threshold corresponds to a subsampled metagenome that still has at least 30-50 ribosomal protein genes sampled. Vertical dashed line denoted the 5% cutoff. Horizontal dashed line denotes a cutoff of 10 ribosomal proteins that is typically used for individual genomes and appears to be too low for metagenomes (30-50 is a better cutoff).

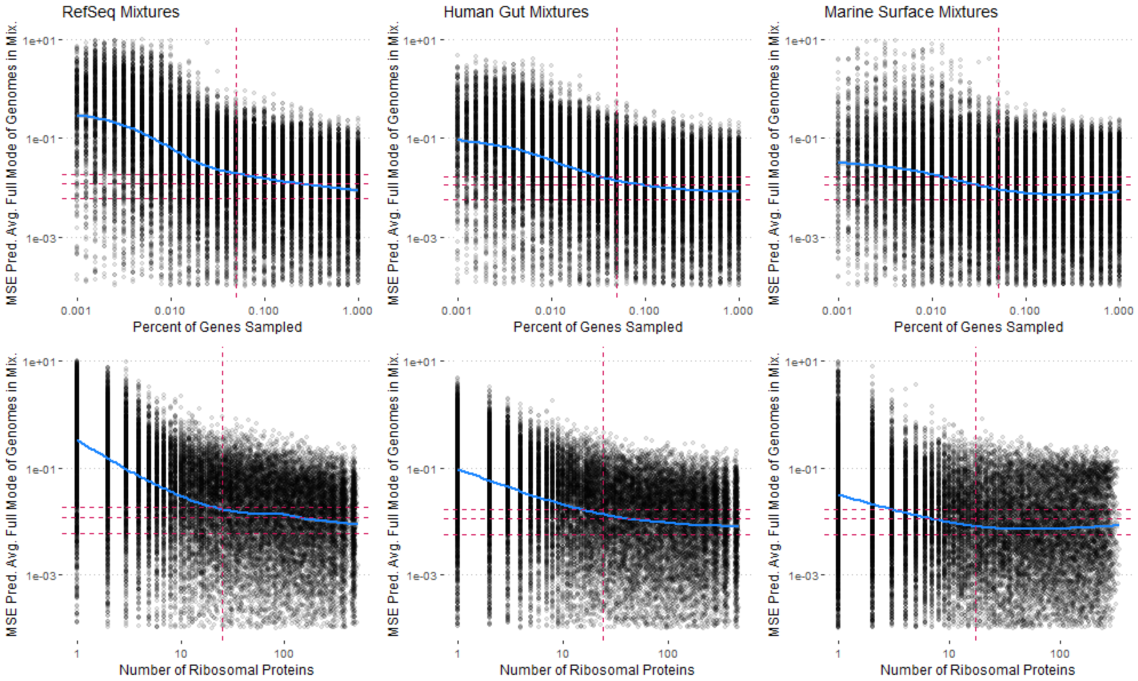

Figure S19: Predicted maximum growth rates (unweighted mode, no coverage information) from genome mixtures are accurate under subsampling as long as at least 5% of genes are sampled. After this threshold errors increase dramatically. Vertical dashed line denoted the 5% cutoff. Horizontal dashed lines denotes mean error of prediction from non-subsampled mixtures and that error value doubled and tripled.

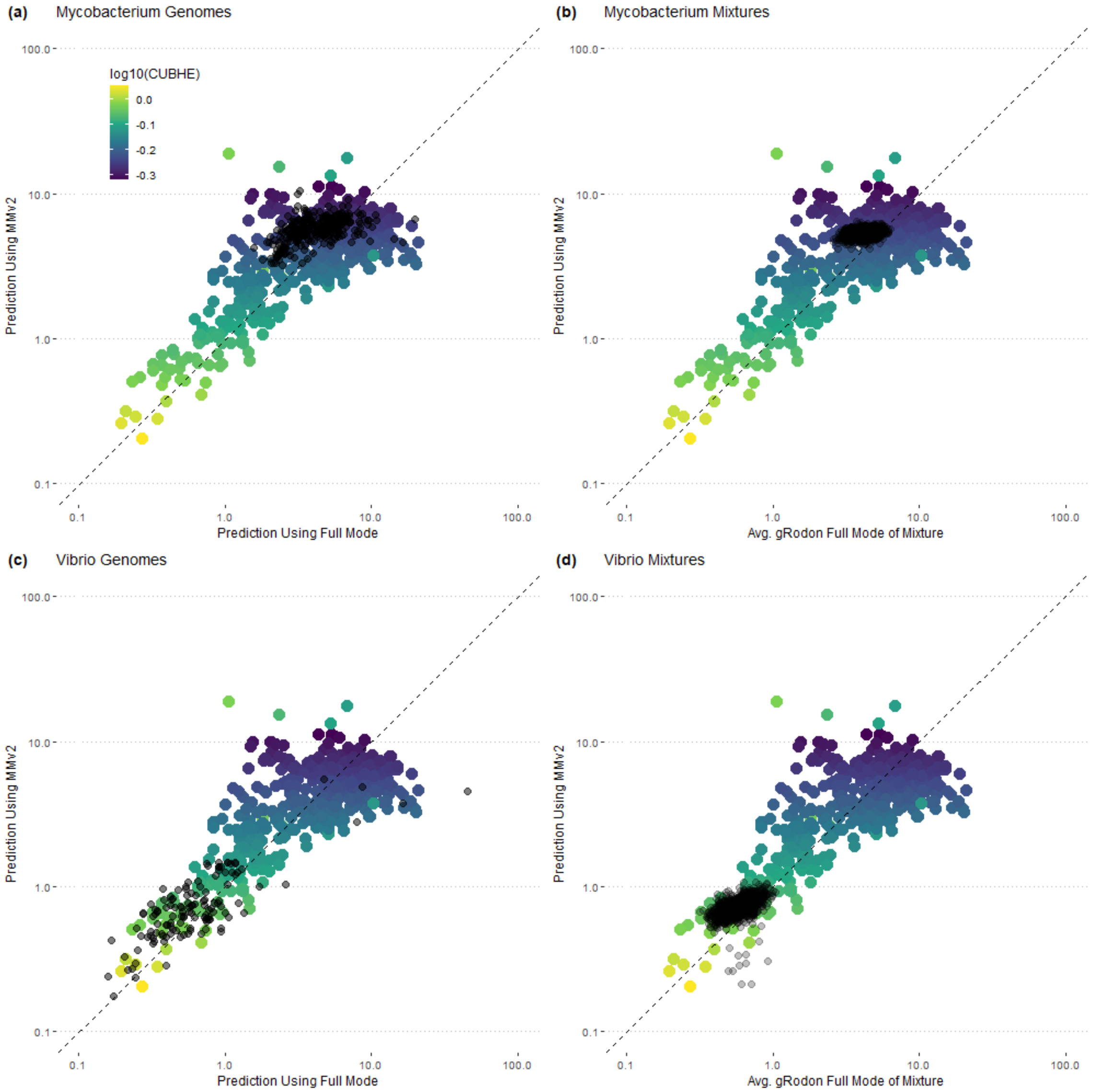

Figure S20: Max. growth rate predictions capture growth rate variation across single-genus species mixtures. (a,c) Variation in growth rate predictions across (a) Mycobacterium or (c) Vibrio species drawn from GTDB207 shown as black points. (b,d) Community-level predictions on 10-species mixtures drawn from either (b) only Mycobacteria, or (d) Vibrio shown as black points. Larger colored-in points correspond to the distribution of predictions of the entire training set used to fit gRodon models, with the color indicating the mean codon usage bias of the highly expressed genes.

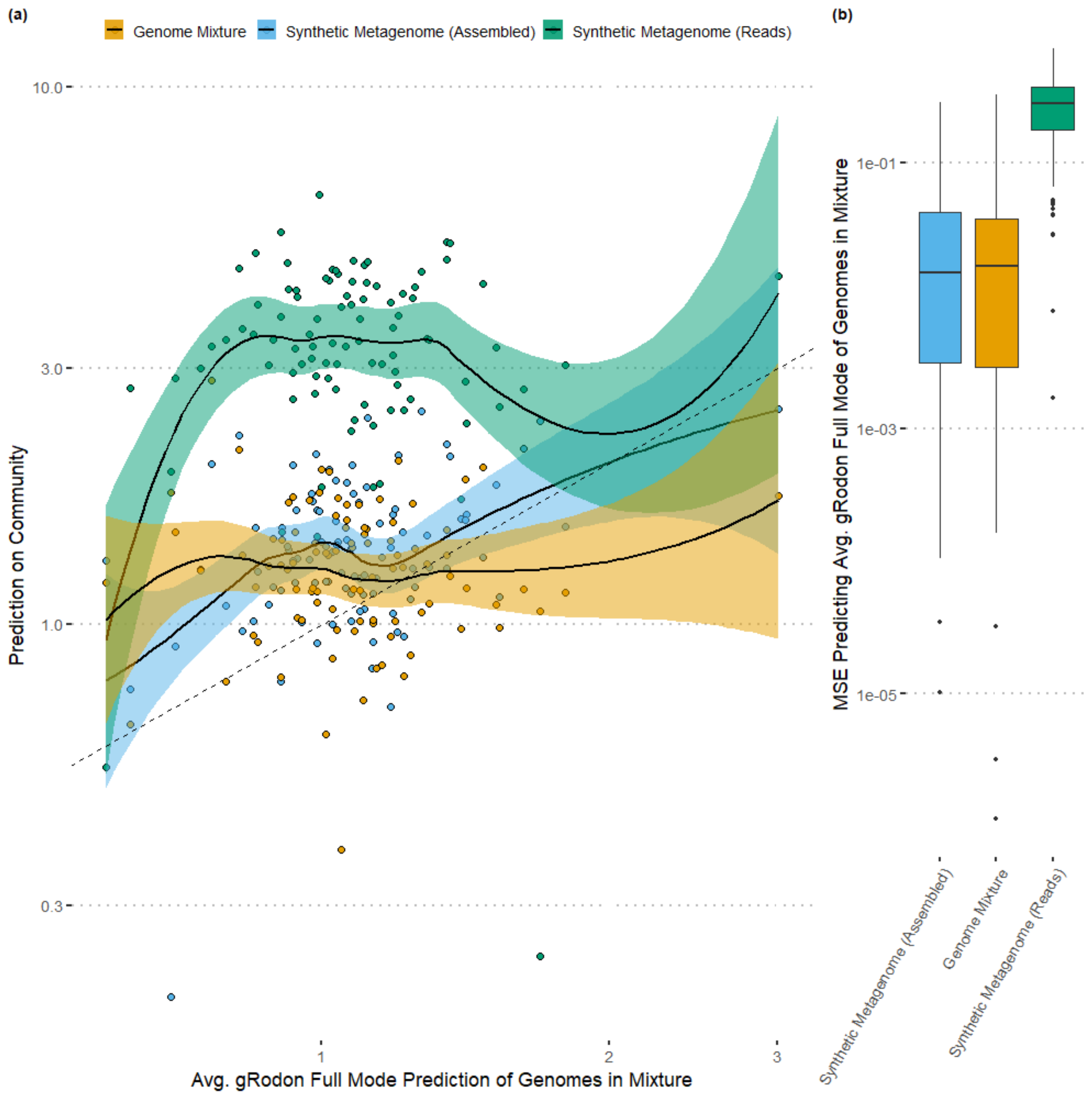

Figure S21: The results of Fig 4 are unchanged when sampling a larger number (10%) of genes predicted from reads for each sample. (a-b) Doubling time predictions on genome mixtures, assembled synthetic metagenomes, and the reads of synthetic metagenomes for the same sets of organisms were compared.
